# ‘*Candidatus* Tisiphia’ is a widespread Rickettsiaceae symbiont in the mosquito *Anopheles plumbeus* (Diptera: Culicidae)

**DOI:** 10.1101/2023.02.27.529723

**Authors:** Helen R. Davison, Jessica Crozier, Stacy Pirro, Doreen Werner, Gregory D.D. Hurst

**Author notes:** **Corresponding Author** Helen R. Davison. **Author contributions** Conceptualisation, supervision, methodology and funding acquisition – HRD and GDDH Resources – HRD, DW, and SP Investigation – JC and HRD Analysis, Visualisation, and Data Curation – HRD Original Draft Preparation – HRD, GDDH. **Data availability statement** Genome bioproject accessions: PRJNA694375 and PRJNA901697. PCR sequences are deposited in accessions OQ512853-OQ512860 Supplementary data is available online at: https://doi.org/10.6084/m9.figshare.21865416.v1.

## Abstract

Symbiotic bacteria alter host biology in numerous ways, including the ability to reproduce or vector disease. Deployment of symbiont control of vector borne disease has focused on *Wolbachia* interactions with *Aedes* and is hampered in *Anopheles* by a lack of compatible symbioses. Previous screening found the symbiont ‘*Ca*. Tisiphia’ in *Anopheles plumbeus*, an aggressive biter and potential secondary vector of malaria parasites and West Nile virus. We screen *An. plumbeus* samples collected over a ten-year period across Germany and use climate databases to assess environmental influence on incidence. We find a 95% infection rate that does not fluctuate with broad environmental factors. Microscopy suggests the infection is maternally inherited based on strong localisation throughout the ovaries. Finally, we assemble a high-quality draft genome of ‘*Ca*. Tisiphia’ to explore its phylogeny and potential metabolism. This strain is closely related to those found in *Culicoides* midges and shows similar patterns of metabolic potential. *An. plumbeus* provides a viable avenue of symbiosis research in anopheline mosquitoes, which to date have one other proven infection of a heritable symbiont. Additionally, it provides future opportunity to study the impacts of ‘*Ca*. Tisiphia’ on natural and transinfected hosts, especially in relation to reproductive fitness and vector efficiency.

## Introduction

Bacterial symbionts in insects form vital components of their host’s biology, ecology, and evolution. They are known to influence how insects reproduce, how they respond to environmental stress, and they interact with pathogens (Dunbar *et al*., 2007; Himler *et al*., 2011; Vega, Arribére and Castro-Vazquez, 2012; Hendry, Hunter and Baltrus, 2014; Xie *et al*., 2014; Hayashi *et al*., 2016). Several species of symbionts are vertically inherited from one generation to the next, usually through the maternal germline, and may become intrinsically linked with their host physiology, metabolism, and development (Buchner, 1965; Zchori-Fein, Borad and Harari, 2006; Moran, McCutcheon and Nakabachi, 2008; Kremer *et al*., 2009; Giorgini *et al*., 2010). Most importantly, symbionts have been deployed in the control of vector populations and vector competence (Hoffmann *et al*., 2011).

Success in symbiont-mediated disease control has been restricted to species from the genus *Aedes*. Transinfection with *Wolbachia* from a drosophilid fly has been successfully used to alter vector competence and lower risk of catching dengue fever from *Aedes aegypti* (Linnaeus, 1762) in endemic areas (Hoffmann *et al*., 2011; Walker *et al*., 2011; Pereira *et al*., 2018). However, important vectors like *Anopheles* are rarely naturally infected with *Wolbachia*, and species within the group are commonly unreceptive to artificial *Wolbachia* infections (Hughes *et al*., 2014). In *Anopheles* mosquitoes, for instance, there is a single well-established case of natural *Wolbachia* infection (Walker *et al*., 2021). Therefore, it is desirable to find potential alternatives that are either more capable of surviving transinfection or alter vector competence in the native host species.

We previously detected the symbiont ‘*Ca*. Tisiphia’ (= Torix group *Rickettsia*) in *Anopheles plumbeus* (Stephens, 1828) in the UK (Pilgrim *et al*., 2021). *An. plumbeus* is broadly distributed across Europe and is an indiscriminate biter. It is also capable of transmitting West Nile virus and malaria parasites, although these pathogens do not natively occur in the majority of the mosquito species’ known distribution range, and vector competence has only been tested in the laboratory setting (Bueno-Marí and Jiménez-Peydró, 2011; Dekoninck *et al*., 2011; Schaffner *et al*., 2012). It has been highlighted as a species that could act as a secondary vector of “tropical” disease agents as changing climate causes their northward spread and their associated primary hosts like *Aedes albopictus* (Skuse, 1894) (Schaffner *et al*., 2012; Heym *et al*., 2017).

‘*Candidatus* Tisiphia’ (= Torix group *Rickettsia*) appear to be particularly associated with hosts deriving from wet or aquatic environments and may originate from symbionts of freshwater ciliates (Driscoll *et al*., 2013; Schrallhammer *et al*., 2013; Kang *et al*., 2014). Infection with this symbiont occur in a broad range of invertebrates, from annelids to gastropods to arthropods (Pilgrim *et al*., 2021), as well as in algae (Hollants *et al*., 2013) and amoebae (Dyková *et al*., 2013). Their relatives in *Rickettsia* are capable of nutritional symbioses, protecting against fungal infections and reproductive manipulation (Hurst *et al*., 1994; Giorgini *et al*., 2010; Hendry, Hunter and Baltrus, 2014; Bodnar *et al*., 2018). However, the known effects of ‘*Ca*. Tisiphia’ are limited to an association with increased host size in *Torix* leeches, and weak impacts on fecundity in *Cimex lectularius* bedbugs (Kikuchi and Fukatsu, 2005; Thongprem *et al*., 2020). There is no observed congruence of host and symbiont phylogeny, indicating that host shifts occur commonly and that long-standing associations with species are rare. External influence such as temperature or natural enemies can also influence the prevalence of symbionts in host populations (Cass *et al*., 2016; Corbin *et al*., 2017; Leclair *et al*., 2017).

Here we use PCR assays to establish the extent of ‘*Ca*. Tisiphia’ infection in *An. plumbeus* mosquitoes across Germany collected by DW and through citizen science initiatives, and assess associations with temperature, precipitation, and forest type. We also sequence and assemble a high-quality draft genome for the symbiont ‘*Ca*. Tisiphia’ and provide evidence of vertical transmission of the symbiont through the maternal germline through FISH imaging. The symbiont genome is examined through bioinformatics approaches to establish potential nutritional or protective symbioses.

## Experimental Procedures

### Collection of *Anopheles plumbeus* (Stephens, 1828)

Two hundred and fifty five *An. plumbeus* specimens were collected from 2012-2021 across Germany by DW and citizen volunteers as part of the mosquito atlas (Mückenatlas) project (Werner *et al*., 2014). These were stored in 70% ethanol or dry (see supplementary materials for storage and exact geographic information). *Post hoc* analysis indicated storage method did not affect detection of symbionts by PCR assay.

Specimens were also collected by HD as larvae and raised to adults in water collected from their larval pools. These specimens were either killed by flash freezing in liquid nitrogen prior to genomic DNA extraction, or in 4% paraformaldehyde solution prior for fluorescence imaging.

### DNA extraction and PCR screening of *Anopheles plumbeus* for *‘Ca*. Tisiphia*’*

Promega Wizard® Genomic DNA Purification kit was used for DNA preparation. DNA quality was then checked with a combination of HCO/C1J primers HCO_2198 (5’-TAA ACT TCA GGG TGA CCA AAA AAT CA-3’)/CIJ_1718 (5’-GGA GGA TTT GGA AAT TGA TTA GT-3’) (Folmer *et al*., 1994; Hajibabaei *et al*., 2005; Siozios *et al*., 2020). *Ca*. Tisiphia presence was assessed with a PCR assay amplifying the 320-bp region of the 17 kDa OMP gene Ri17kD_F (5’-TCTGGCATGAATAAACAAGG-3’)/Ri17kD_R (5’-ACTCACGACAATATTGCCC-3’) (Pilgrim *et al*., 2017). PCR conditions used were as follows: 95 °C for 5 min, followed by 35 cycles of denaturation (94 °C, 30 s), annealing (54 °C, 30 s) and extension (72 °C, 120 s).

A selection of ‘*Ca*. Tisiphia’ amplicons from positive samples across time and space were Sanger-sequenced through Eurofins barcode service and identity confirmed by comparing the sequences to the NCBI database via BLAST homology searches. These sequences are deposited in accessions OQ512853-OQ512860.

### Association of symbiont prevalence with geographic and climatic information

Annual average monthly temperature and precipitation data were retrieved for each sample’s coordinate and year from TerraClim (Abatzoglou *et al*., 2018) which has a spatial resolution of ∼4-km (1/24th degree). Forest cover data was retrieved from Copernicus land datasets for 2018 (European Union, 2018) and raster data for forest type extracted in QGIS 3.16 (QGIS.org, 2020) within a 3km radius of each sample location. *Anopheles plumbeus* has historically been recorded to have a maximum flight range of up to 13km (Becker *et al*., 2010). However, this is based on one single study from 1925 and is not verified by other sources. As such we chose an estimated range of 3km based on the average flight ranges other anopheline mosquito species (Becker *et al*., 2010; Verdonschot & Besse-Lototskaya, 2014). Scikitlearn’s standard scaler (Pedregosa *et al*., 2011) was applied to data before performing a generalised linear model with a binomial logit link function on data with the following formula:

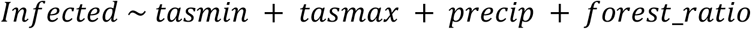

All statistics and geographic inferences were carried out in Python with the packages Statsmodel and Scikit-learn (Rossum and Drake, 2009; Seabold and Perktold, 2010; Pedregosa *et al*., 2011). QGIS 3.16 was used to produce maps and extract raster data for forest types before passing it to Python for analysis (QGIS.org, 2020). All other figures were produced with Matplotlib and Seaborn (Hunter, 2007; Waskom and Seaborn development team, 2020).

### Fluorescence in situ microscopy (FISH)

Reproductive organs of a single female and a single male were dissected and incubated in cold 4% paraformaldehyde for 3 hours, agitated gently every 30 minutes, then washed with cold PBS for 5 minutes two times. Tissue was stained with Hoescht 33342 (that binds double-stranded DNA) for 30 minutes at room temperature, then hybridised overnight at room temperature with hybridisation buffer (5X SSC, 0.01% SDS, 30% formamide) and 5′-CCATCATCCCCTACTACA-(ATTO 633)-3′ oligonucleotide probe specific to ‘*Ca*. Tisiphia’ 16S rRNA (Pilgrim *et al*., 2017). Hybridised tissue was washed in wash buffer (5X SSC, 0.01% SDS) at 48°C for 60 minutes with gentle shaking every 20 minutes. Samples were then mounted in Vectashield. Images were taken with a ZEIS LSM 880 confocal microscope through ZEIS Zen black, and final images were annotated in Inkscape Ver 1.2 (Inkscape Project, 2020).

### De novo sequencing, assembly, and annotation

A combination of short and long read sequencing was used to construct scaffolds for the ‘*Ca*. Tisiphia’ genome. For short reads, Iridian Genomes extracted and processed DNA of one male for Illumina sequencing deposited under bioproject accession PRJNA694375. The short reads of *An. plumbeus* were cleaned with Trimmomatic 0.36 (Bolger, Lohse and Usadel, 2014) and quality checked with FASTQ (Babraham Bioinformatics, 2019). For long reads, genomic DNA from one male was extracted with Qiagen Genomic-tip for ultra-low PacBio sequencing carried out by the Centre for Genomic Research, University of Liverpool. Long read sequences are deposited under bioproject accession number PRJNA901697.

A combination of long and short reads were used to assemble as complete a genome for ‘*Ca*. Tisiphia’ as possible. First, the ‘*Ca*. Tisiphia’ genome was identified in the Illumina short reads and assembled through Minmap2, MEGAHIT and MetaBAT2 as per the pipeline used in Davison *et al*. (2022). Second, PacBio HiFi long read sequences were assembled using Flye 2.9. 1-b1780 with the ‘-meta’ flag to improve sensitivity for low coverage reads. Third, the long read assembly was queried against a local blast database of *Rickettsia* and ‘*Ca*. Tisiphia’ genomes (including the Illumina assembly from the first step) to identify sequences belonging to this strain of ‘*Ca*. Tisiphia’. Lastly, the long read assembly was polished with the Illumina reads using Polypolish v0.5.0 (Wick and Holt, 2022) with default settings to give 23 final scaffolds.

### Phylogeny and metabolic predictions

Annotation of the final assembly was carried out with InterProScan v5 (Jones *et al*., 2014). Metabolic pathway prediction for presence and completion was carried out through Anvi’o 7 using KEGG kofams and COG20 (Aramaki *et al*., 2020; Eren *et al*., 2021; Galperin *et al*., 2021). NRPS pathways were investigated with AntiSMASH 6.0 (Blin *et al*., 2021).

The ‘*Ca*. Tisiphia’ strain for *An. plumbeus* was compared to the other existing ‘*Ca*. Tisiphia’ genomes through Anvi’o 7. A core genome consisting of 205 gene clusters that contain a total of 3690 genes was found through Anvio-7. Phylogenies were estimated from single copy gene clusters with IQTREE 2.2.0.3 using Model Finder Plus and with 1000 ultrafast bootstraps (Kalyaanamoorthy *et al*., 2017; Hoang *et al*., 2018; Minh *et al*., 2020). The model selected by Model Finder Plus is Q.plant+F+R4. A supporting phylogeny to confirm the identity of 17 kDa OMP genes was produced with the model TIM2+F.

## Results and Discussion

### Distribution and predicted environment

*‘Ca*. Tisiphia’ was observed to infect *An. plumbeus* across all sites examined in Germany, with 95% specimens positive on PCR assay. The few negative specimens were found to mostly occur in the southeast of the country (Figure 1 and Supplementary Figure 1). The infection seems to be stable, and there is no evidence of frequency change over time, with samples from all years spanning 2012 to 2022 displaying similar rates of infection (Supplementary Figure 1, Supplementary Table S1, Supplementary Figure 2).

**Figure 1.**
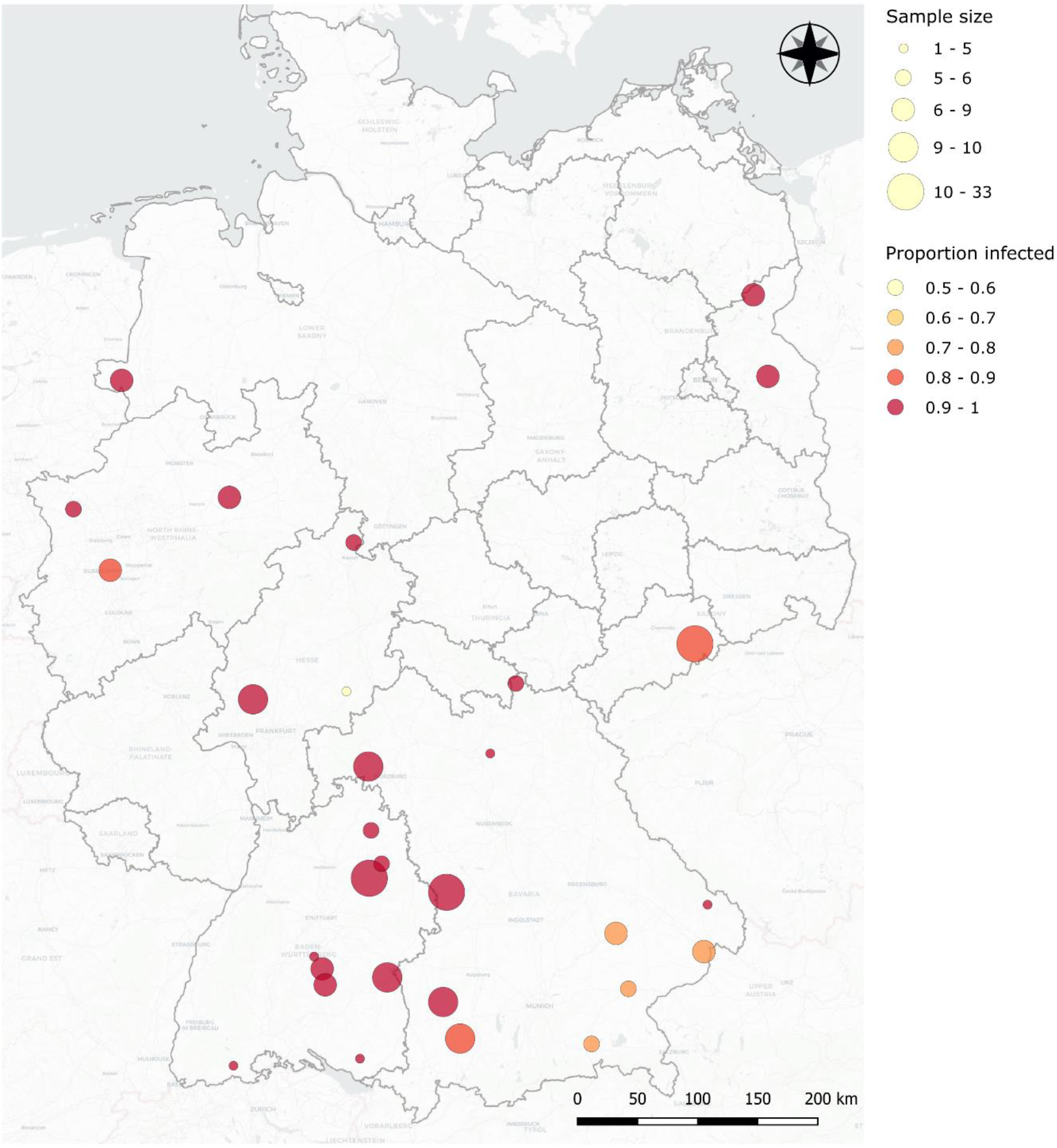
**Map of ‘*Ca*. Tisiphia’ infection rates across Germany** where the size of the circle represents the number of individuals sampled and the colour indicates the proportion of ‘*Ca*. Tisiphia’ infected individuals. Red = 90-100% infection to light yellow = 50-60% infection. Source data can be found in Supplementary Table S1.

There is no clear evidence of an influence on ‘*Ca*. Tisiphia’ infection rates in *An. plumbeus* caused by average minimum or maximum temperature, precipitation or forest types (Figure 2 and Supplementary Figure 2 and 3). While there appears to be a significant effect of precipitation on the number of uninfected individuals, this could be an artifact of increased water availability leading to more mosquitoes and thus a higher chance of detecting rarer uninfected individuals (Supplementary Figure 2). No variation is unsurprising as it appears to be a very high prevalence infection. We also acknowledge that using climate databases to retroactively find data is not as accurate as field measurements. However, results agree with previous field observations of *Rickettsia* infection in *Acyrthosiphon pisum* in Japan (Tsuchida *et al*., 2002). We chose to use the high resolution TerraClim database, but this may still mask small differences in microenvironments are limited to mostly abiotic data. We encourage future symbiosis research to consider environmental measurements to describe the ecology of these organisms more comprehensively.

**Figure 2.**
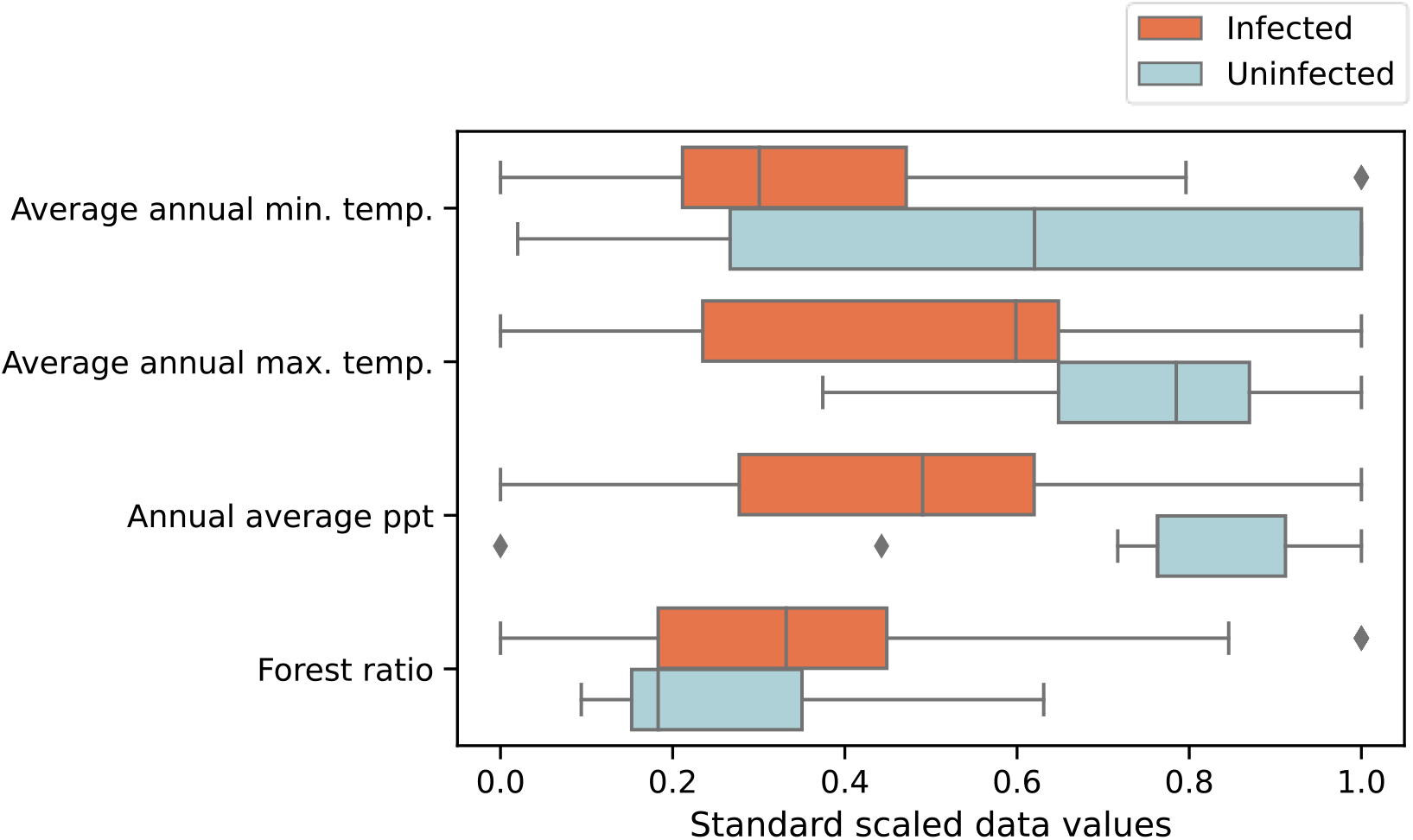
**Standardised and scaled environmental data** comparing Uninfected (N=13) and Infected (N=237) by environmental variable. Source data can be found in Supplementary Table S1.

#### Phylogeny and metabolism

The bacteria sequenced from *An. plumbeus* is most closely related to a ‘*Ca*. Tisiphia’ found in the biting midge *Culicoides newsteadi* (Figure 3). All infections tested are the same strain of ‘*Ca*. Tisiphia’ (Supplementary Figure 4). General features of both genomes are consistent with other ‘*Ca*. Tisiphia’ (Table 1 and Supplementary Data S2-S4); TsAplum has a single full set of rRNAs (16S, 5S and 23S), and GC content is ∼33%. It also has several repeat domains (Table 1) which are associated with protein-protein interactions and are prevalent in *Wolbachia* symbionts (Siozios *et al*., 2013; Rice, Sheehan and Newton, 2017).

**Table 1.**
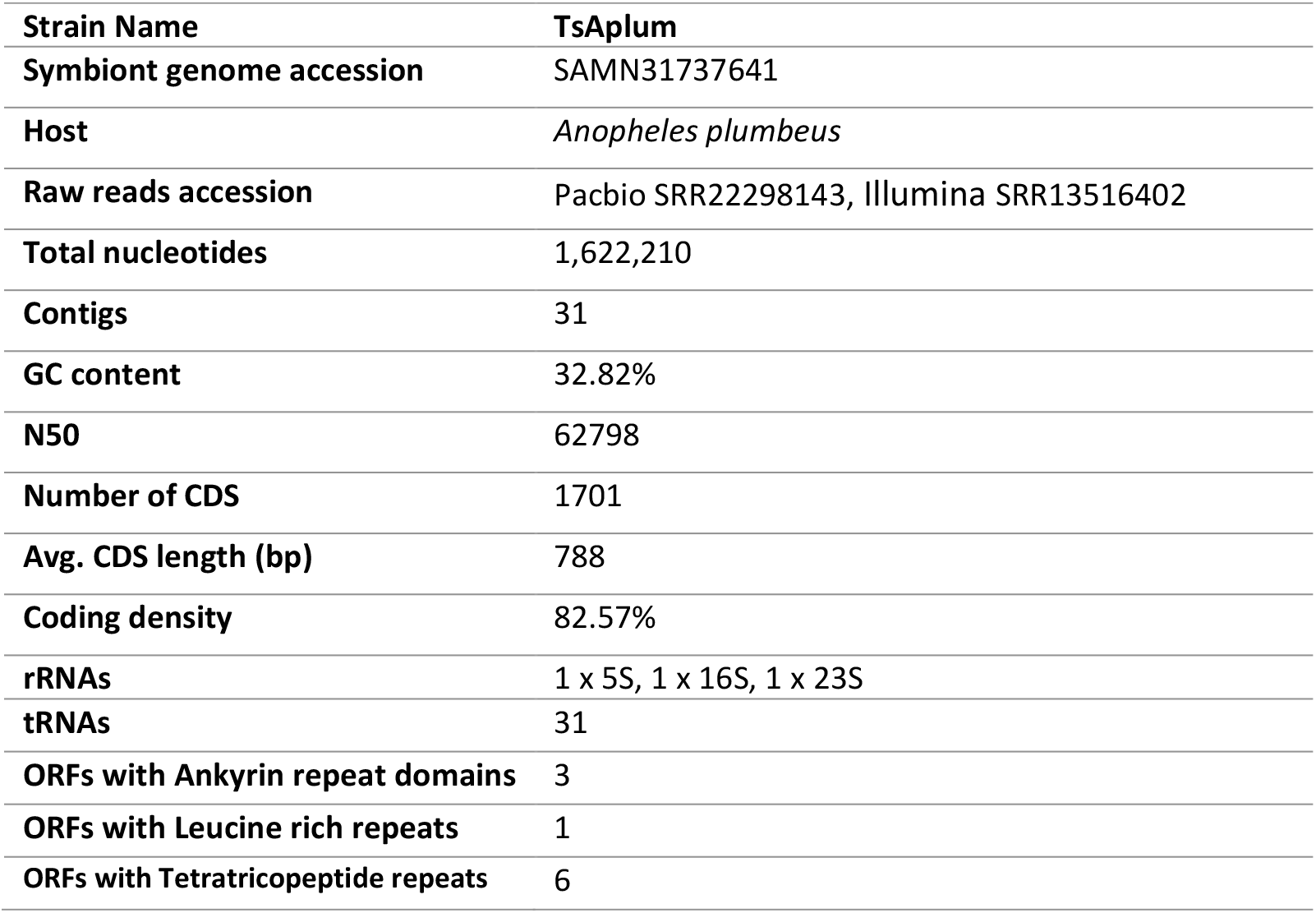
Summary of the genome assembly for TsAplum.

**Figure 3.**
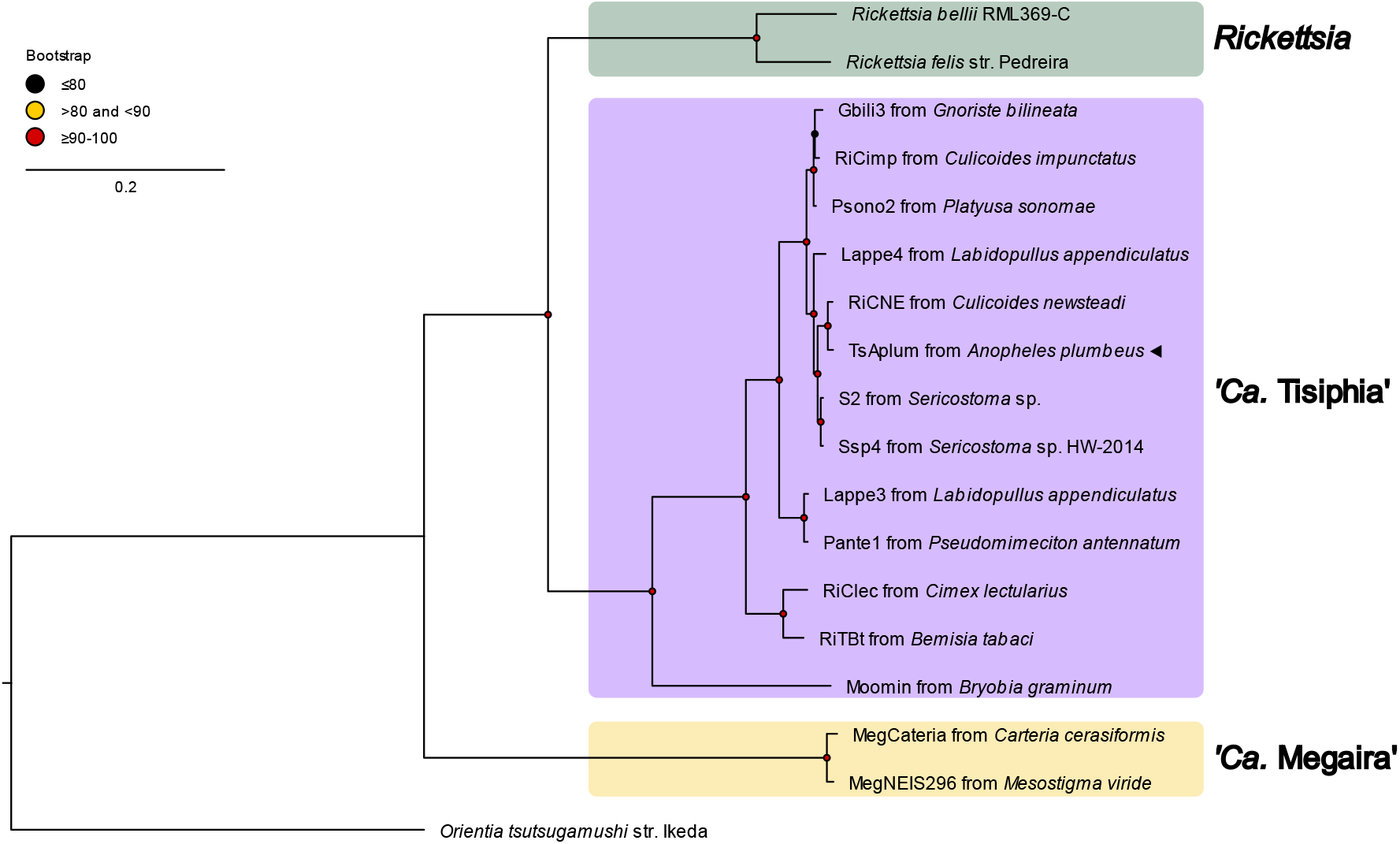
Genome wide phylogeny of ‘*Ca*. Tisiphia’ and ‘*Ca*. Megaira’. Maximum likelihood (ML) phylogeny constructed from 205 single copy gene clusters that contain a total of 3690 genes. New genomes are indicated by ◂ and bootstrap values based on 1000 replicates are indicated with coloured circles (red = 91-100, yellow = 81-90, black <= 80). Accession numbers for each genome are available in Supplementary Table S2.

Overall ‘*Ca*. Tisiphia’ found in *An. plumbeus* mirrors the metabolic potential found in other members of its genus (Figure 4 and Figure 5). It does not have any obvious metabolic pathway that would contribute to nutritional symbiosis such as B vitamin production nor any NRPS/PKS system for small molecule synthesis (Supplementary Tables S3 and S4). It does have several toxin/anti-toxin systems as well as secretion pathways Tat, Sec, VirB (Type IV), all of which are essential in various symbiont-host interactions (Masui, Sasaki and Ishikawa, 2000; Meloni *et al*., 2003; Wu *et al*., 2004; Dale and Moran, 2006; Tseng, Tyler and Setubal, 2009). Additionally, it has a number of ORFs containing ankyrin- and leucine-rich repeats which are thought to be important in interactions with cognate eukaryotic proteins (Siozios *et al*., 2013; Rice, Sheehan and Newton, 2017). Thus, the genome itself, whilst firmly placing the symbiont in the context of the genus and identifying relatedness to other strains, does not raise obvious hypotheses about the impact of infection on the host. Phenotype studies are required to properly assess the influence of this bacteria on its host. Key studies would address the factors driving the spread of the symbiont into the population (testing for beneficial aspects of infection, cytoplasmic incompatibility, and paternal inheritance) and impacts on viral infection and transmission outcomes.

**Figure 4.**
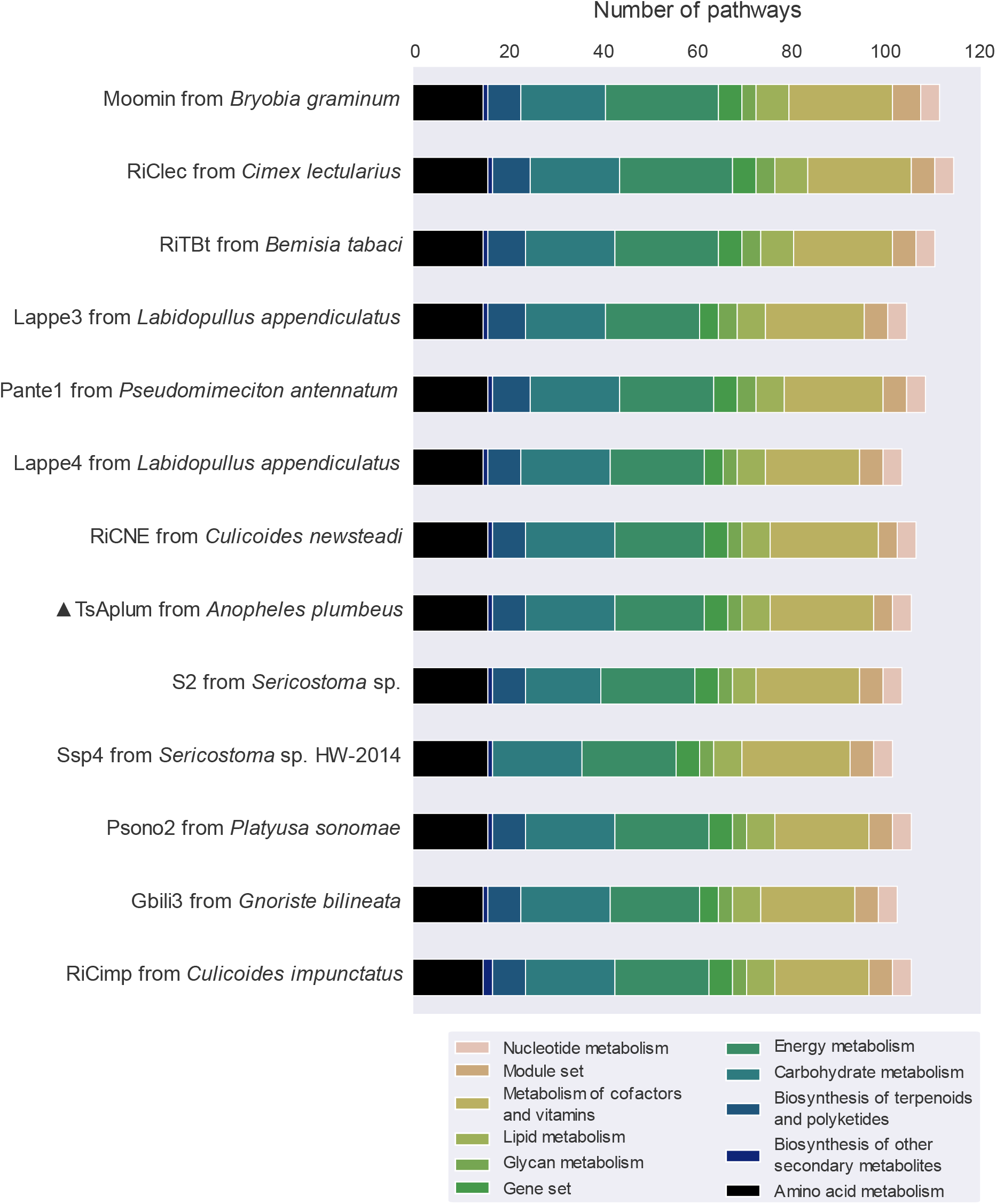
KEGG module distribution in ‘*Ca*. Tisiphia’. The number of pathways found per genome annotated by KEGG module category for ‘*Ca*. Tisiphia’. Full metadata can be found in Supplementary Data 1. ▴ indicates the genome assembled in this study. Full metadata can be found in Supplementary Tables S3 and S4.

**Figure 5.**
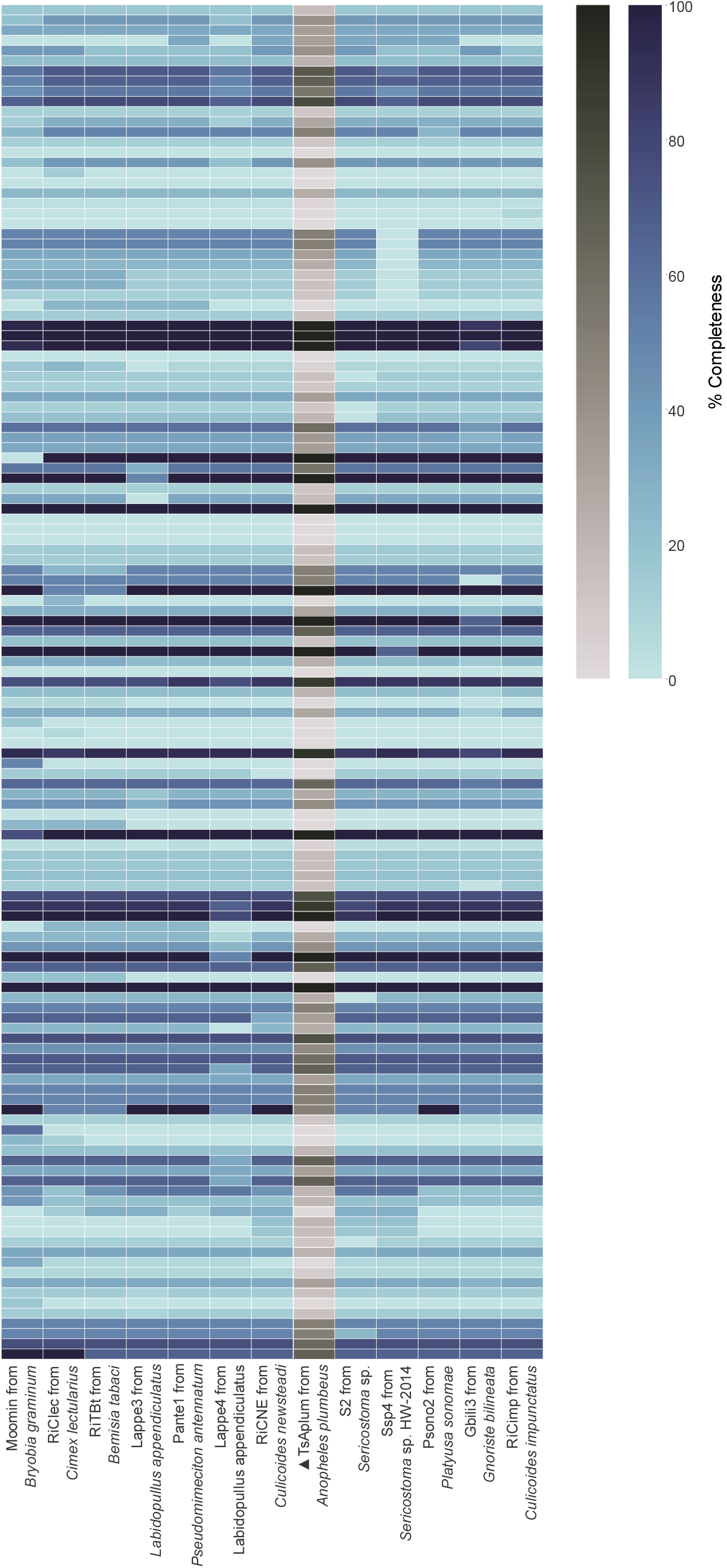
Predicted completeness of KEGG kofam metabolic pathways across ‘Ca. Tisiphia’. The genome assembled in this chapter is coloured grey and indicated with▴ Full metadata can be found in Supplementary Tables S3 and S4.

#### FISH imaging

*‘Ca*. Tisiphia’ is observed in oocytes and oviduct branches but was not detected in testes (Figure 6 versus Supplementary Figure 5). Localisation and clear polarity of the infection in ovaries strongly suggest that this is a maternally inherited infection (Figure 6). The bacteria cluster around the oocyte of the primary follicles as well as in the lateral ducts and secondary follicles. In *Drosophila melanogaster, Wolbachia* is similarly polarised to one end of the primary follicles to the oocyte (Ferree *et al*., 2005), and in *Proechinophthirus fluctus*, their endosymbionts *Sodalis* appears to use the lateral oviducts to access the ovaries (Boyd *et al*., 2016).

**Figure 6.**
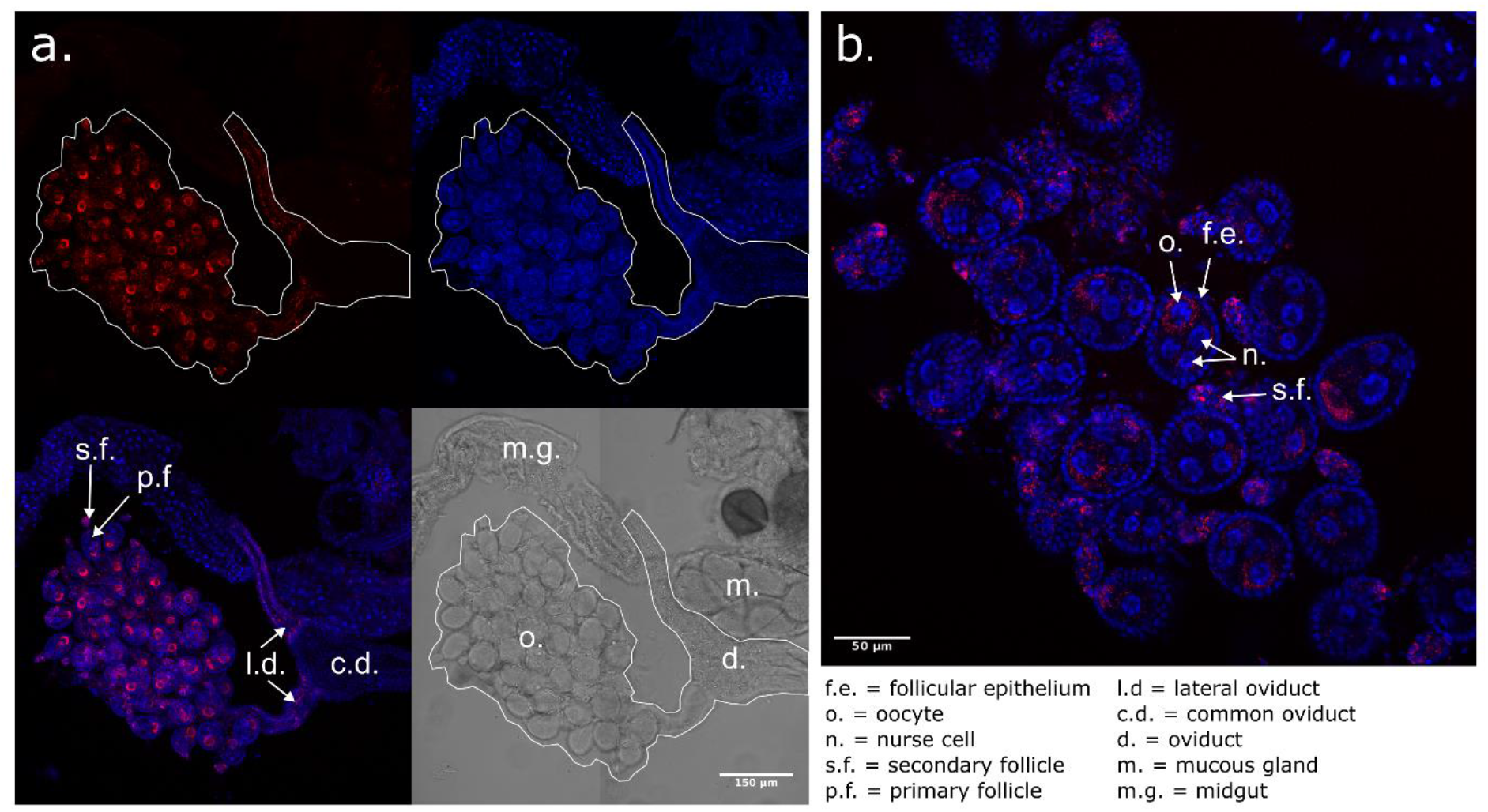
Fluorescence in situ microscopy images of *Anopheles plumbeus* ovaries infected with ‘*Ca*. Tisiphia’. Red shows ‘*Ca*. Tisiphia’ stained with ATTO-633, blue are host nuclei stained with Hoechst. Panels show a) the whole female reproductive organ outlined in white and a breakdown of each light channel and b) a close up of the ovaries showing localised infection within the primary and secondary follicles. White bars indicate a) 150 micrometres and b) 50 micrometres.

#### Final conclusions

*Anopheles plumbeus* and its ‘*Ca*. Tisiphia’ make a good potential model for symbioses in *Anopheles* mosquitoes as well as for ‘*Ca*. Tisiphia’ infection more generally. Outside of *An. plumbeus*, there is only a single well substantiated case of *Wolbachia* and no other symbiont in anopheline mosquitoes (Walker *et al*., 2021). The infection in *An. plumbeus* is clearly evidenced, likely heritable, and occurs in a species that has seen previous success as a laboratory colony (Kotter, 2005). Beyond this, *An. plumbeus* is a species of interest with the capability to carry West Nile virus and *Plasmodium* parasites (Dekoninck *et al*., 2011; Schaffner *et al*., 2012). ‘*Ca*. Tisiphia’ in *An. plumbeus* provides a viable avenue for symbiont-mediated vector modification and control to be tested in anopheline species. It is also an example of a temporally and spatially stable infection of non-pathogenic Rickettsiaceae and a good foil to fluctuating systems like Belli *Rickettsia* in *Bemisia tabaci* (Bockoven *et al*., 2020).

Future work should also establish the effects of this symbiont in transinfection in alternative hosts alongside the native *An. plumbeus* host. Other symbionts like *Wolbachia* are known to produce functionally interesting phenotypes related to vector competence when transferred from the original host into other, naïve species (Moreira *et al*., 2009). Alongside this, impacts on host function and physiology, and potential means of spread into natural populations would need to be assessed. A first step to establishing transinfection would be to isolate the ‘*Ca*. Tisiphia’ infection into cell culture, which would also represent an important community resource for onward study.

In summary, ‘*Ca*. Tisiphia’ is found in 95% of *An. plumbeus* individuals from Germany and forms a well-established, stable, and heritable infection that persists across space and time. Metabolic potential is typical of similar symbiotic bacteria species, and we find no evidence of large-scale environmental factors influencing rates of infection. However, ‘*Ca*. Tisiphia’ and *An. plumbeus* provide a unique opportunity to study the effects of a native symbiont infection in anopheline mosquitoes, as well as explore its potential use for disease mitigation in other species that cannot be infected with currently used symbionts.

## Ethical approval statement

All samples were collected in accordance with ethical standards for insect collection.

## Supplementary Figures

**Supplementary Figure 1.**
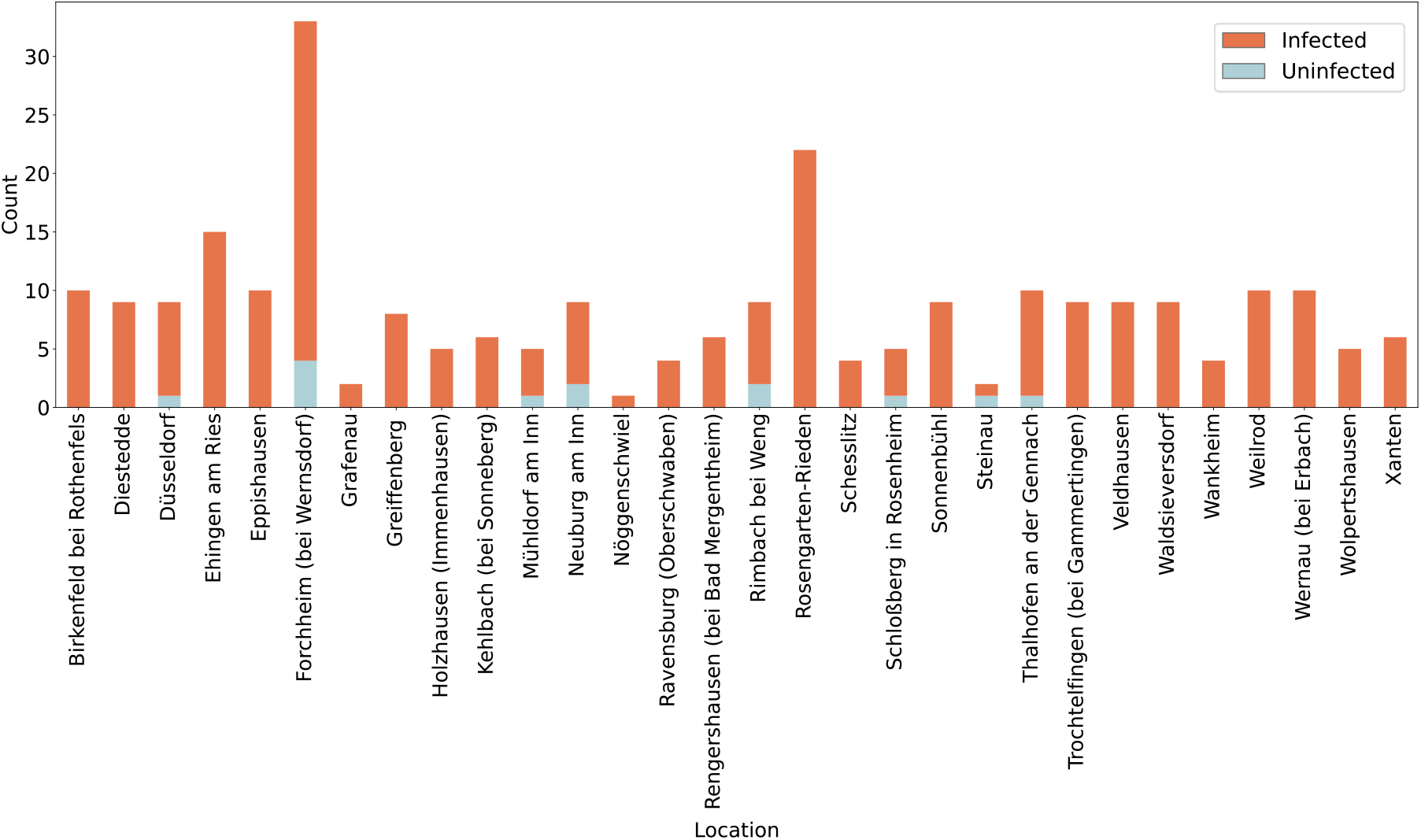
‘*Ca*. Tisiphia’ infection rates by site’. Infected samples are shown in orange, uninfected are shown in light blue. Source data can be found in Supplementary Table S1.

**Supplementary Figure 2.**
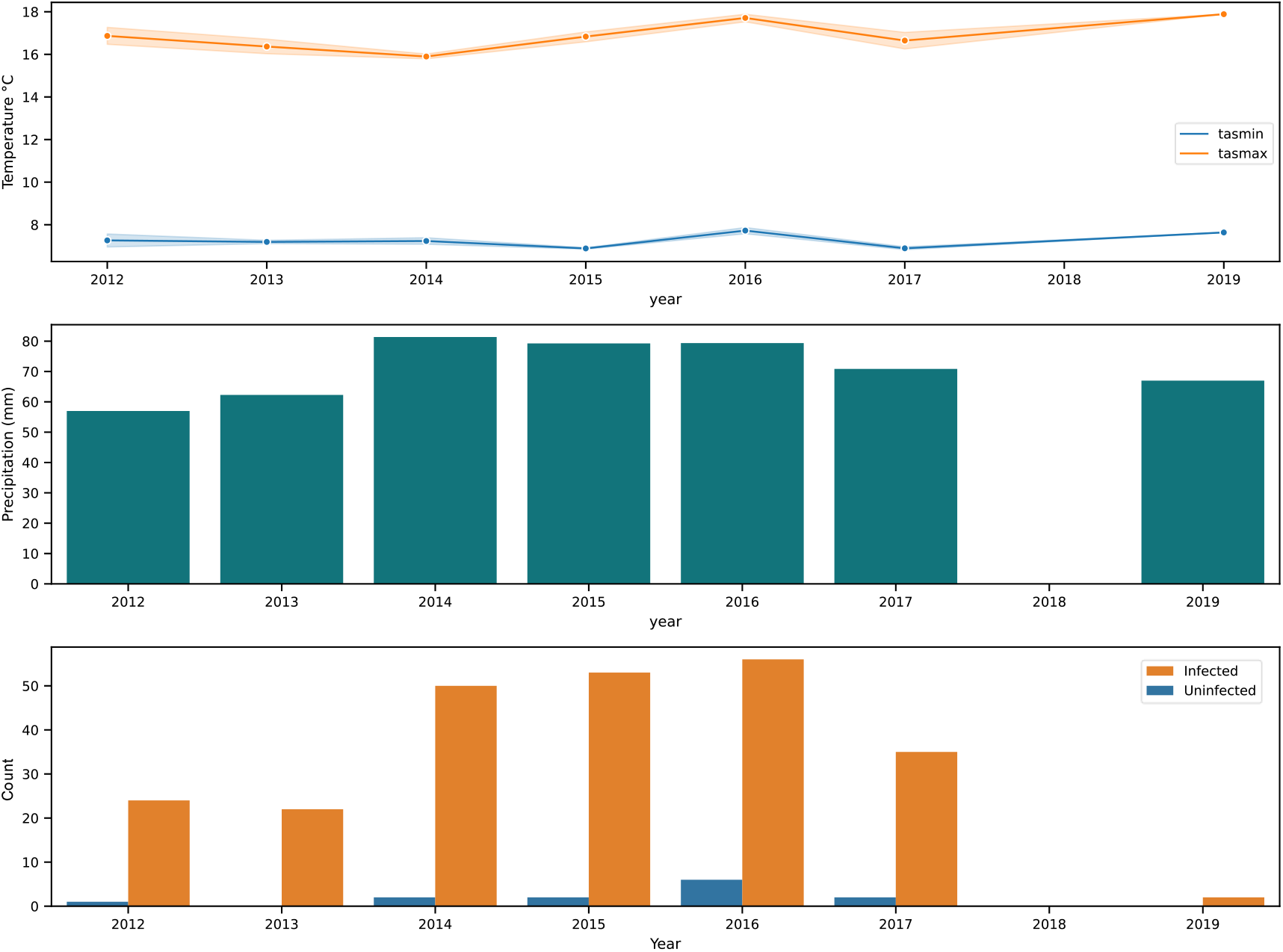
Environmental data for *Anopheles plumbeus* collection sites across Germany extracted from the TerraClim database. (Top) average annual minimum and maximum temperature across all *An. plumbeus* collection sites in Germany. (Middle) average annual precipitation across all sites. (Bottom) counts of infected and uninfected individuals across all sites where dark blue = infected and orange = uninfected.

**Supplementary Figure 3.**
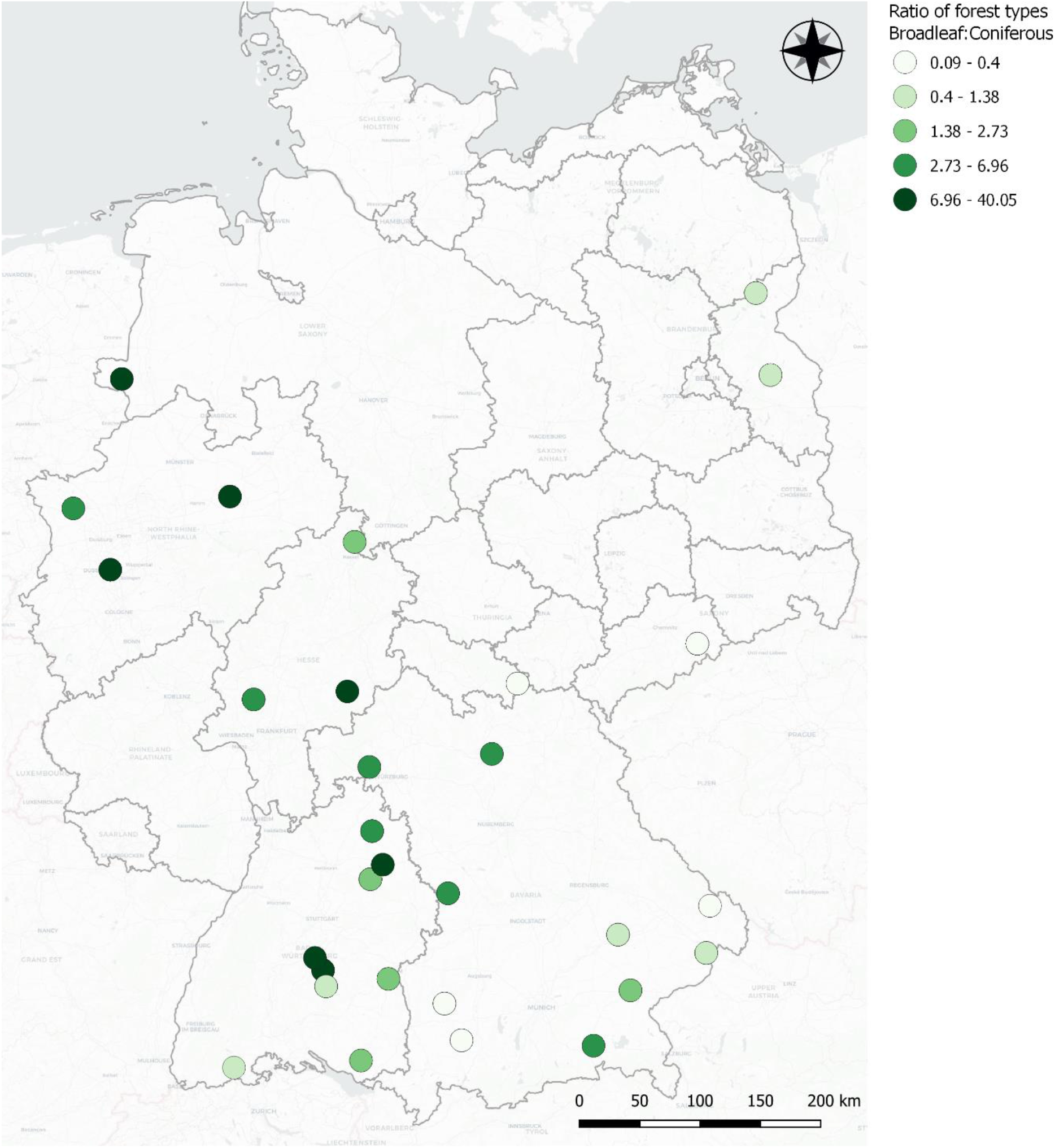
The ratio of broadleaf to coniferous forest in a 3km radius of each collection site. Darker green indicates more broadleaf, lighter green indicates closer to equal proportions. Source data can be found in Supplementary Table S1.

**Supplementary Figure 4.**
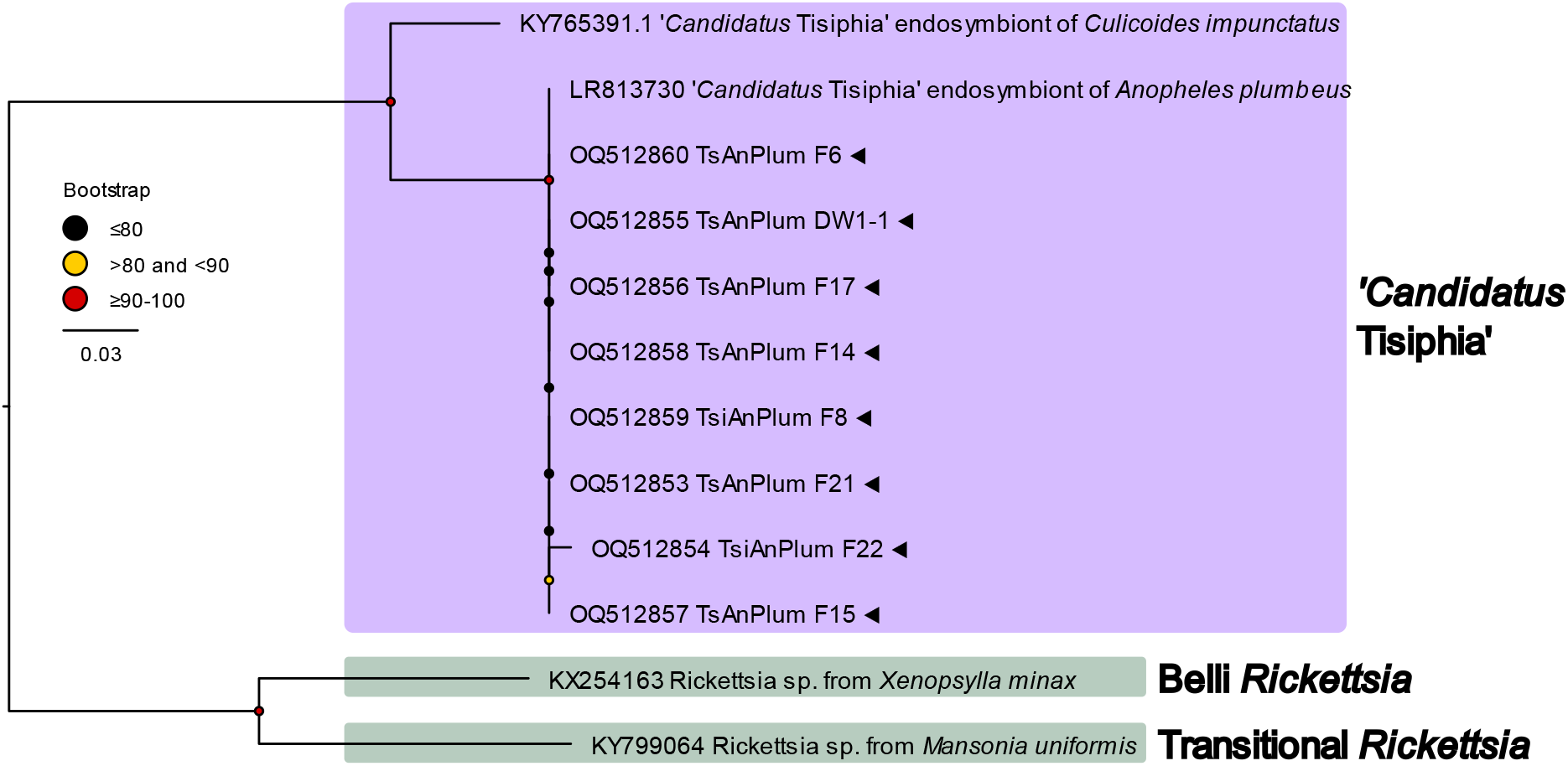
Maximum likelihood tree for 17 kDa surface antigen (omp) for ‘*Candidatus* Tisiphia*’* extracted from *An. plumbeus*. New genomes are indicated by ◂ and bootstrap values based on 1000 replicates are indicated with coloured circles (red = 91-100, yellow = 81-90, black <= 80).

**Supplementary Figure 5.**
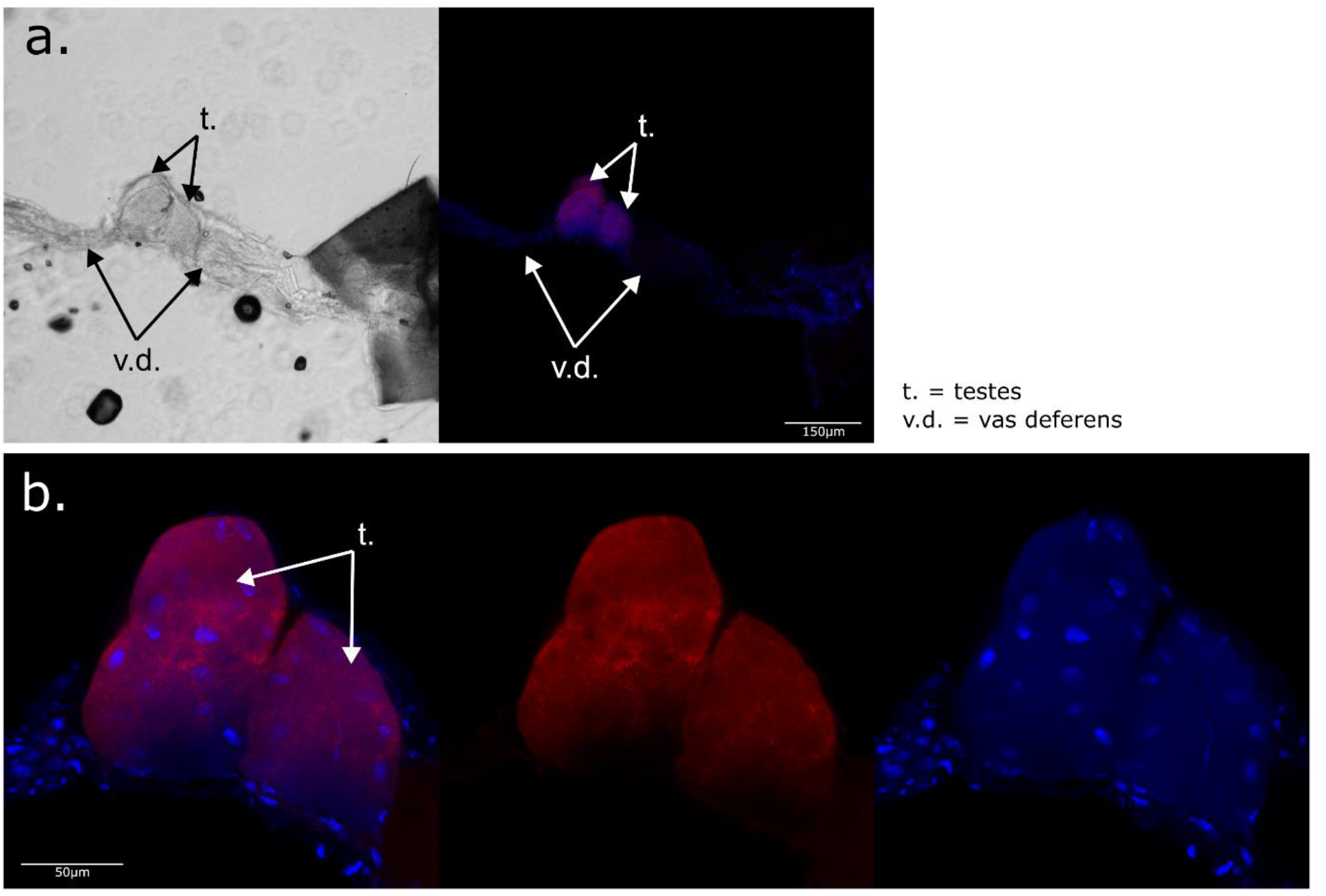
Fluorescence in situ microscopy images of *Anopheles plumbeus* testes. Blue are host nuclei stained with Hoechst 33342, Red is ATTO-633 auto-fluorescence in the testes not ‘*Ca*. Tisiphia’ staining. White bars indicate a) 150 micrometres and b) 50 micrometres.

